# Bovine coronavirus evolution preserves spike architecture while remodeling key functional regions

**DOI:** 10.64898/2026.08.05.743018

**Authors:** Haneen Tarabih, Jimmy Asiku, Yael Levi-Kalisman, Moriya Slavin, Ran Zalk, Anat Shahar, Jenna Weisz, Rina Fraenkel, Dan David, Nir Kalisman, Asaf Sol, Netanel Tzarum

## Abstract

Bovine coronavirus (BCoV) is a major pathogen of cattle and the closest known ancestor of the human coronavirus OC43, yet how viral evolution reshapes spike protein structure and function remains poorly understood. Here, we combined comparative genomics, glycoproteomics, cryo-EM, and antigenic characterization to define the structural mechanisms underlying spike evolution across representative BCoV lineages. We identify a previously unrecognized lineage-specific N-glycosylation site in contemporary European viruses and validate its occupancy by glycoproteomics. High-resolution cryo- EM structures reveal that BCoV evolution preserves the overall prefusion architecture of the spike glycoprotein while selectively remodeling key functional regions involved in receptor recognition, conformational dynamics, and antigenicity. Comparative analysis with OC43 demonstrates increased conformational heterogeneity of the receptor-binding loop, whereas the Mebus vaccine strain exhibits enhanced membrane-proximal stalk flexibility despite maintaining thermal stability. Finally, structural modeling together with antibody-binding experiments reveals substantial antigenic remodeling despite >90% spike sequence identity between BCoV and OC43. Together, these findings establish a mechanistic framework for Embecovirus spike evolution and provide structural insights that may inform the development of vaccines based on contemporary circulating strains.

## Introduction

Bovine coronavirus (BCoV) is an enveloped, positive-sense RNA virus belonging to the genus *Betacoronavirus*, subgenus *Embecovirus*, first isolated in 1972 (Stair *et al*, 1972; Vlasova & Saif, 2021), and is a major viral pathogen affecting cattle worldwide. BCoV causes substantial economic losses to the dairy and beef industries through reduced milk production, decreased weight gain, treatment costs, and mortality (Boileau & Kapil, 2010). Outbreaks can affect cattle of all ages and impose high direct and indirect costs, including production losses, veterinary care, and disease control (Decaro *et al*, 2008; Saif, 2010).

BCoV is associated with three major clinical syndromes in cattle. Winter dysentery (WD) is an acute, highly contagious enteric disease affecting adult cattle, characterized by hemorrhagic diarrhea, reduced milk production, and substantial production losses (Tsunemitsu & Saif, 1995). In young calves, BCoV is a major cause of neonatal calf diarrhea (CD), which is associated with severe enteritis and increased mortality (Tsunemitsu & Saif, 1995). BCoV also contributes to respiratory disease in cattle of all ages as a component of the bovine respiratory disease complex (BRDC), one of the most economically significant infectious diseases of the cattle industry (Saif, 2010). The ability of closely related BCoV strains to infect both the intestinal and respiratory tracts highlights the remarkable biological plasticity of the virus and raises important questions regarding the molecular determinants of tissue tropism and host adaptation.

The BCoV genome encodes five structural proteins, including the spike (S) glycoprotein and the hemagglutinin-esterase (HE), a hallmark of *Embecoviruses*. The trimeric spike mediates viral attachment and membrane fusion and is therefore the principal determinant of host-cell entry (Popova & Zhang, 2002). Unlike SARS-CoV, SARS-CoV-2, and MERS-CoV, which utilize protein receptors, BCoV recognizes 9-*O*-acetylated sialic acids through its N-terminal domain (NTD) (Peng *et al*, 2012). As the major surface glycoprotein exposed on the virion, the spike is also the primary target of neutralizing antibodies (nAbs).

Among the *Embecoviruses*, human coronavirus OC43 (OC43) is the closest evolutionary and structural relative of BCoV (Vijgen *et al*, 2005; Vlasova & Saif, 2021). Phylogenetic analyses suggest that OC43 emerged following the zoonotic transmission of BCoV into humans approximately 130 years ago (Vijgen *et al*., 2005). Consistent with this shared ancestry, both viruses recognize 9-*O*-acetylated sialic acids and exhibit highly similar spike architectures (Peng *et al*., 2012). High-resolution cryo-EM structures of the OC43 spike have provided detailed insights into receptor recognition and antigenic epitopes (Bangaru *et al*, 2022; Tortorici *et al*, 2019; Wang *et al*, 2022), establishing a structural framework for investigating how BCoV spike proteins have evolved.

Like other CoVs, BCoV evolves through mutation and recombination, reshaping both the amino acid sequence and glycosylation landscape of its spike glycoprotein (Franzo *et al*, 2020; Vlasova & Saif, 2021). N-linked glycans contribute not only to immune evasion but also to protein folding, conformational stability, receptor accessibility, and spike dynamics (Allen *et al*, 2023; Watanabe *et al*, 2020; Yang *et al*, 2025). Likewise, sequence variation within the receptor-binding NTD can modify spike structure and antigenicity. Despite remarkable advances in cryo-EM and glycoproteomic analyses of human coronaviruses over the past decade, comparable structural information for BCoV remains scarce. Consequently, it is unknown whether the receptor-binding architecture, glycosylation landscape, and conformational dynamics observed in OC43 are conserved in BCoV or have diverged during the evolution of contemporary bovine viruses.

Commercial BCoV vaccines are based primarily on historical isolates, most notably the prototypic Mebus strain, which was isolated more than five decades ago (Mebus *et al*, 1972). However, BCoV has continued to evolve, raising concerns that historical vaccine strains may no longer accurately represent the structural and antigenic properties of contemporary circulating viruses. Indeed, recent molecular epidemiological studies from multiple geographic regions have shown that contemporary BCoV strains have diverged genetically from historical isolates (David *et al*, 2021; Saif, 2010; Suzuki *et al*, 2020; Zhu *et al*, 2022), the structural and molecular consequences of this evolutionary divergence remain unknown. In particular, it is unclear how lineage-specific mutations remodel spike glycosylation, receptor-binding architecture, conformational dynamics, and antigenic properties.

Here, we investigated the structural mechanisms by which contemporary BCoV evolution has reshaped the spike glycoprotein. Comparative genomic analyses identified a previously unrecognized lineage-specific N-linked glycosylation site that is conserved among contemporary European viruses and confirmed its occupancy by glycoproteomic. We then determined high-resolution cryo-EM structures of spike proteins representing historical and contemporary BCoV lineages and integrated these structural data with complementary biophysical and antigenic analyses. Together, our findings reveal that ongoing BCoV evolution remodels spike architecture through coordinated changes in glycosylation, conformational dynamics, and antigenicity, providing a mechanistic framework for understanding *Embecovirus* evolution and informing the rational design of vaccines based on contemporary circulating strains.

## Results

### Comparative phylogenetic analysis identifies a lineage-specific glycosylation site in contemporary BCoV spike proteins

To determine how ongoing BCoV evolution has reshaped the spike glycoprotein, we first examined the genetic diversity of contemporary BCoV isolates circulating in Israel by sequencing the complete spike and HE genes from samples collected between 2017 and 2022. Complete spike sequences were obtained from 18 animals originating from 15 dairy farms, whereas complete HE sequences were obtained from 25 animals representing 21 farms. The sampled animals presented with calf diarrhea, winter dysentery, pneumoenteric disease, and respiratory infection (Fig. S1A).

Phylogenetic analysis of the full-length spike sequences showed that the Israeli BCoV isolates segregated into two closely related lineages, with no apparent association between phylogenetic clustering and clinical syndrome, year of isolation, or geographic origin (Fig. 1A). A similar clustering pattern was observed for the HE sequences, indicating that both viral surface glycoproteins evolved in parallel within the Israeli BCoV population (Fig. S1B).

**Figure 1.**
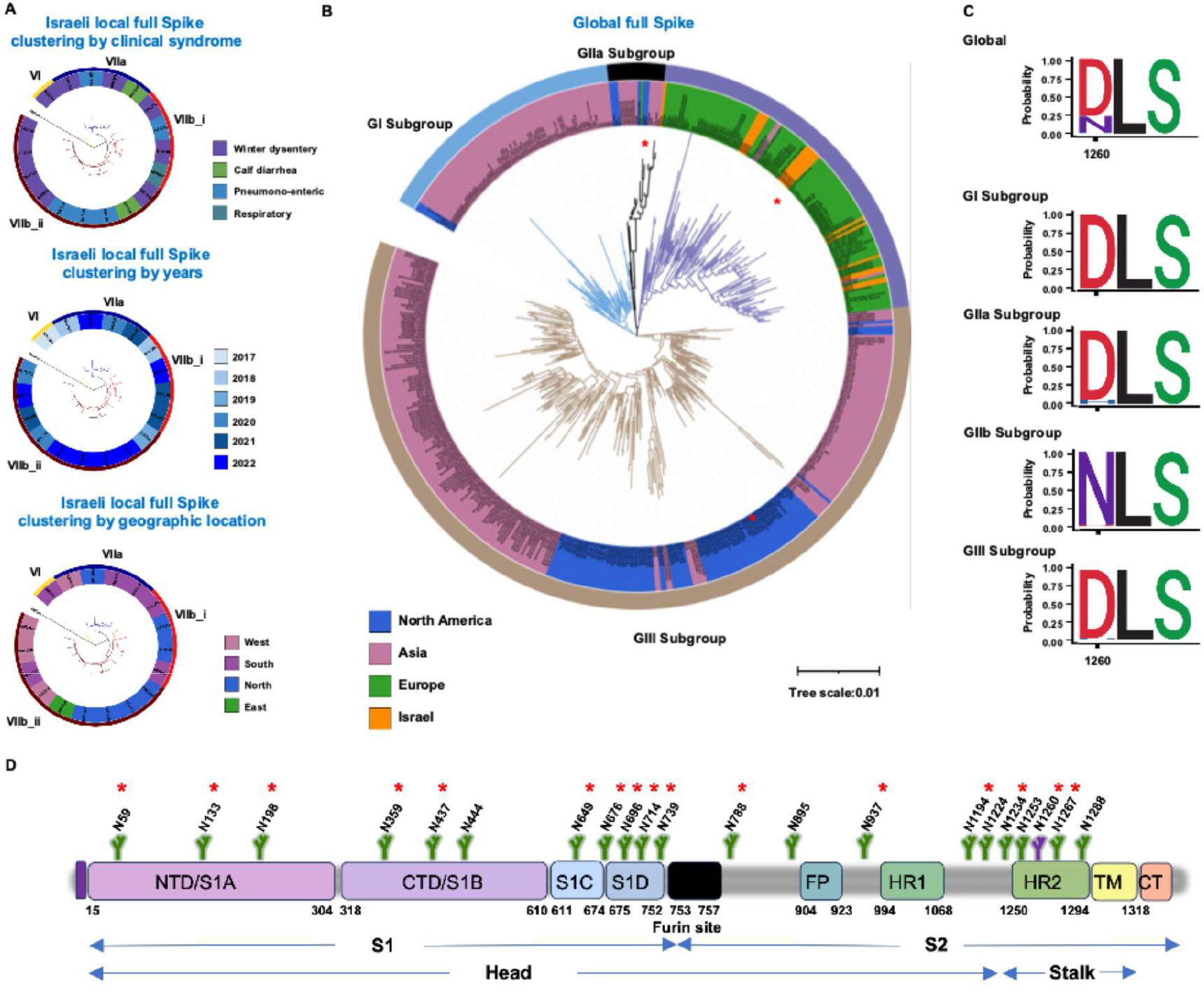
Phylogenetic and sequence analyses identify a lineage-specific N1260 glycosylation site in the BCoV spike glycoprotein. **(A)** Phylogenetic analysis of full-length spike glycoprotein sequences from Israeli BCoV isolates collected between 2017 and 2022. Isolates are annotated according to year of isolation, geographic location, and clinical syndrome. **(B)** Maximum-likelihood phylogenetic tree of full-length BCoV spike sequences from Israeli isolates together with representative global strains. Four major phylogenetic lineages (GI, GIIa, GIIb, and GIII) are identified. The three strains selected for structural and biochemical characterization in this study (Mebus, ISR2182, and BRCV2014) are indicated by red asterisks. **(C)** Sequence logo analysis of residue 1260 in the BCoV spike glycoprotein. The upper panel shows the amino acid distribution across all available BCoV spike sequences, whereas the lower panels show the corresponding distributions for the GI, GIIa, GIIb, and GIII phylogenetic lineages. **(D)** Schematic representation of the BCoV spike glycoprotein showing the domain organization and predicted N-linked glycosylation sites. The lineage-specific N1260 glycosylation site is highlighted in purple. Glycosylation sites experimentally confirmed by LC-MS/MS are indicated by red asterisks.

To place the Israeli isolates in the context of global BCoV diversity, we constructed a phylogenetic tree using all publicly available full-length spike and HE sequences from the NCBI database (Fig. 1B, Fig. S1C and Supplementary Table 1). The global phylogeny resolved four major lineages (GI, GIIa, GIIb, and GIII). All Israeli isolates clustered within the contemporary European GIIb lineage, whereas the prototype Mebus strain grouped within GIIa. This evolutionary framework informed the selection of three representative strains spanning the major phylogenetic groups (Mebus, ISR2182, and BRCV2014) for subsequent structural and biochemical analyses.

Sequence logo analysis of the spike proteins identified six lineage-defining amino acid positions (Supplementary results and Fig. S1D). Among these, the GIIb-specific D1260N substitution was unique in generating a canonical N-linked glycosylation sequon (N-X-S/T), thereby distinguishing contemporary European and Israeli strains from the remaining BCoV lineages (Fig. 1C).

Analysis of the predicted glycosylation landscape revealed that N1260 represented the only variable N-linked glycosylation site among the major BCoV lineages, whereas all other predicted glycosylation sites were conserved (Fig. S1E). Given the strong conservation and functional importance of S2 glycans among *Betacoronaviruses* (Allen *et al*., 2023; Watanabe *et al*., 2020) (Fig. S1F), we next asked whether the newly acquired N1260 sequon is occupied in the mature spike glycoprotein.

### Experimental validation of the lineage-specific N1260 glycosylation site

To experimentally validate occupancy of the lineage-specific N1260 glycosylation site, we generated a prefusion-stabilized recombinant ISR2182 spike ectodomain using established coronavirus spike engineering strategies (Esposito *et al*, 2020; Schaub *et al*, 2021). The construct design, purification, and biochemical characterization are described in the Supplementary Results (Fig. S1G-H). The purified spike ectodomain was treated with Endoglycosidase H (EndoH) prior to subsequent trypsin or chymotrypsin digestion and LC-MS/MS analysis (Fig. S1I). Endo H treatment simplifies glycan heterogeneity while retaining a single N-acetylglucosamine residue on glycosylated asparagine residues, enabling confident assignment of occupied glycosylation sites.

Complementary digestion with trypsin and chymotrypsin generated extensive sequence coverage across the spike ectodomain, including 20 out of the 21 potential glycosylation sites (excluding position 1124, Fig. S1J-K). LC-MS/MS analysis confirmed N-linked glycosylation at 16 of the 20 predicted recovered sites, indicating that the recombinant spike protein largely recapitulates the expected glycosylation profile (Fig. 1D, S1E). Importantly, the peptide containing residue N1260 was identified with the characteristic residual GlcNAc modification in both trypsin- and chymotrypsin-treated samples, confirming that the lineage-specific N1260 is occupied by N-linked glycosylation in the mature spike protein. Moreover, we did not identify any peptide in which the N1260 GlcNAc modification was missing, indicating that this is a particularly strong glycosylation site with close to 100% occupancy. Together, these findings demonstrate that contemporary BCoV evolution has generated a previously unrecognized lineage-specific N-linked glycosylation site, establishing the first structural signature distinguishing contemporary European viruses from historical BCoV lineages.

### Cryo-EM structures reveal structural differences among representative BCoV lineages

Having identified a lineage-specific glycosylation site in contemporary BCoV strains, we next asked whether ongoing viral evolution has also remodeled the overall architecture of the spike glycoprotein. To address this question, we determined high-resolution cryo-EM structures of prefusion-stabilized spike ectodomains from three representative phylogenetic lineages: the prototypic Mebus strain (GIIa), the contemporary European ISR2182 strain (GIIb), and the contemporary North American BRCV2014 strain (GIII) (Fig. 2A, Fig. S2A). Inclusion of both contemporary lineages enabled us to distinguish lineage-specific structural adaptations from changes associated with the Mebus vaccine strain. Prefusion-stabilized spike ectodomains were transiently expressed and purified as described above (Fig. S2B). The structures were determined at overall resolutions of 2.66 Å for ISR2182, 2.50 Å for Mebus, and 2.52 Å for BRCV2014 (Fig. 2A and Fig. S2C, D), enabling detailed comparative structural analyses.

**Figure 2.**
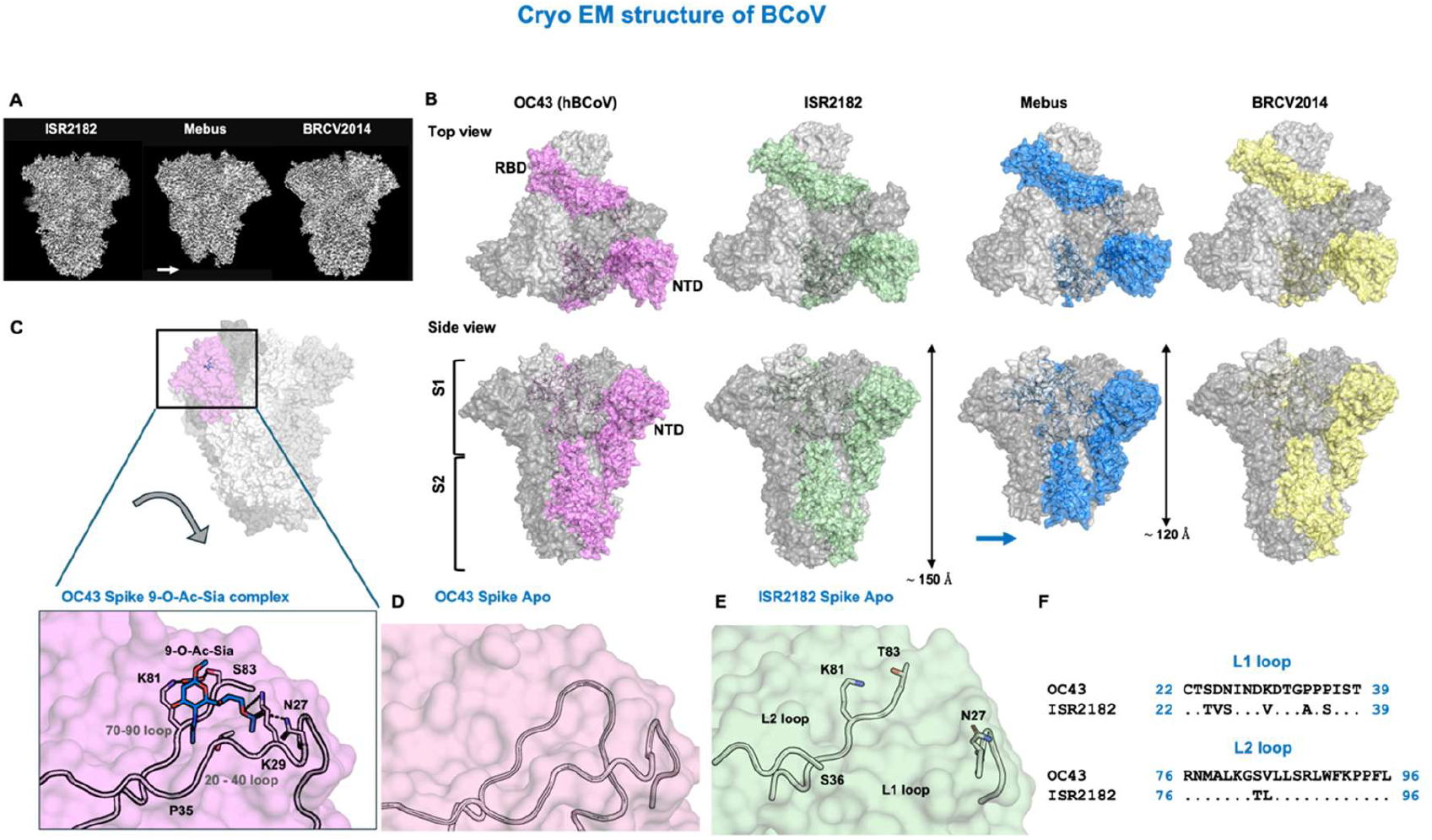
Comparative cryo-EM analysis of representative bovine coronavirus spike glycoproteins. **(A)** Cryo-EM density maps of the prefusion-stabilized spike trimers from the ISR2182 (GIIb), Mebus (GIIa), and BRCV2014 (GIII) strains. **(B)** Structural comparison of the BCoV spike trimers and the previously reported OC43 spike structure (PDB:6NZK). Orthogonal side and top views are shown for each structure. One protomer is colored while the remaining two protomers are shown in light and dark gray. **(C)** Structure of the OC43 spike N-terminal domain (NTD) in complex with 9-*O*- acetylated sialic acid receptor (PDB:6OHW). The receptor-binding pocket is formed primarily by the L1 and L2 loops. Residues involved in receptor recognition are shown as sticks. **(D)** Apo structure of the OC43 NTD showing a similar conformation of the receptor-binding loops in the absence of ligand. **(E)** Apo structure of the ISR2182 NTD. Residues corresponding to the L1 loop are not resolved because of the absence of well-defined cryo-EM density, whereas the L2 loop remains well resolved. **(F)** Sequence alignment of the receptor-binding L1 and L2 loops from OC43 and ISR2182 as representative of BCoV.

Structural analysis of the BCoV Ecto structures indicated that the three spike trimers adopted the canonical prefusion coronavirus architecture, with three S1 subunits surrounding the central trimeric S2 fusion core (Fig. 2B). Most domains were well resolved, whereas the membrane-proximal region of S2, including residue N1260, remained unresolved in all three structures, precluding direct visualization of the lineage-specific glycan. Despite their evolutionary divergence, the ISR2182 and BRCV2014 spike proteins exhibited highly similar overall architectures and closely resembled the previously reported OC43 spike. In contrast, the prototypic Mebus spike displayed markedly reduced density within the membrane-proximal stalk, indicative of increased conformational heterogeneity, while maintaining the same overall prefusion organization. Overall, comparative cryo-EM analysis demonstrates that the BCoV spike has retained a highly conserved prefusion architecture despite ongoing evolution, with structural divergence concentrated in discrete functional regions of the glycoprotein. Furthermore, the overall organization of the glycan shield remained conserved across the three BCoV lineages, despite extensive sequence divergence, whereas structural comparison with OC43 revealed an expanded CTD while preserving its overall fold (Supplementary Result and S2E–G).

### The BCoV receptor-binding L1 loop exhibits conserved conformational flexibility

To determine whether evolution has remodeled the receptor-binding surface of the BCoV spike, we next examined the N-terminal domain (NTD), which mediates receptor recognition through binding to 9-O-Ac-Sia. Previous structural studies of OC43 demonstrated that the receptor-binding pocket is primarily formed by the L1 and L2 loops, with the L1 loop contributing key residues that stabilize the ligand-binding cavity (Fig. 2C) (Tortorici *et al*., 2019). Comparison of apo and receptor-bound OC43 structures showed that these loops adopt nearly identical conformations, indicating that the receptor-binding site is preformed and structurally rigid (Fig. 2C, D). In contrast, residues 28-35 of the corresponding BCoV L1 loop could not be modeled in any of the three spike structures because of the absence of well-defined cryo-EM density, indicating pronounced conformational flexibility within this receptor-binding loop (Fig. 2E and Fig. S2H). Notably, this increased flexibility was observed in all three BCoV spike structures despite their phylogenetic divergence, suggesting that it is a conserved structural feature of BCoV rather than a lineage-specific property. Sequence comparison revealed that the corresponding OC43 L1 loop contains three consecutive proline residues (Pro34-Pro36), whereas the BCoV loop lacks two of these residues (Fig. 2F). Given the well-established role of proline residues in restricting backbone flexibility, these substitutions may contribute to the increased conformational mobility of the BCoV receptor-binding loop. Together, these findings identify increased conformational flexibility of the receptor-binding L1 loop as a conserved structural feature of the BCoV spike, distinguishing it from the closely related OC43. This structural plasticity may have important implications for receptor recognition and antigenicity, which remain to be determined.

### The membrane-proximal stalk of the Mebus spike exhibits increased conformational heterogeneity

Although the three BCoV spike proteins exhibited highly similar overall prefusion architectures, the membrane-proximal stalk of the Mebus strain lacked well-defined cryo-EM density, suggesting greater conformational heterogeneity than in ISR2182 and BRCV2014. Interestingly, this did not affect the rest of the structure, as the Mebus reconstruction achieved the highest overall resolution (2.50 Å). Moreover, the unresolved density does not correspond to a single continuous segment of the primary sequence but instead comprises residues originating from multiple discontinuous regions that converge to form the membrane-proximal stalk in the folded structure (Fig. 3A). These observations indicate increased mobility of the stalk as a structural unit rather than localized flexibility of an individual loop.

**Figure 3.**
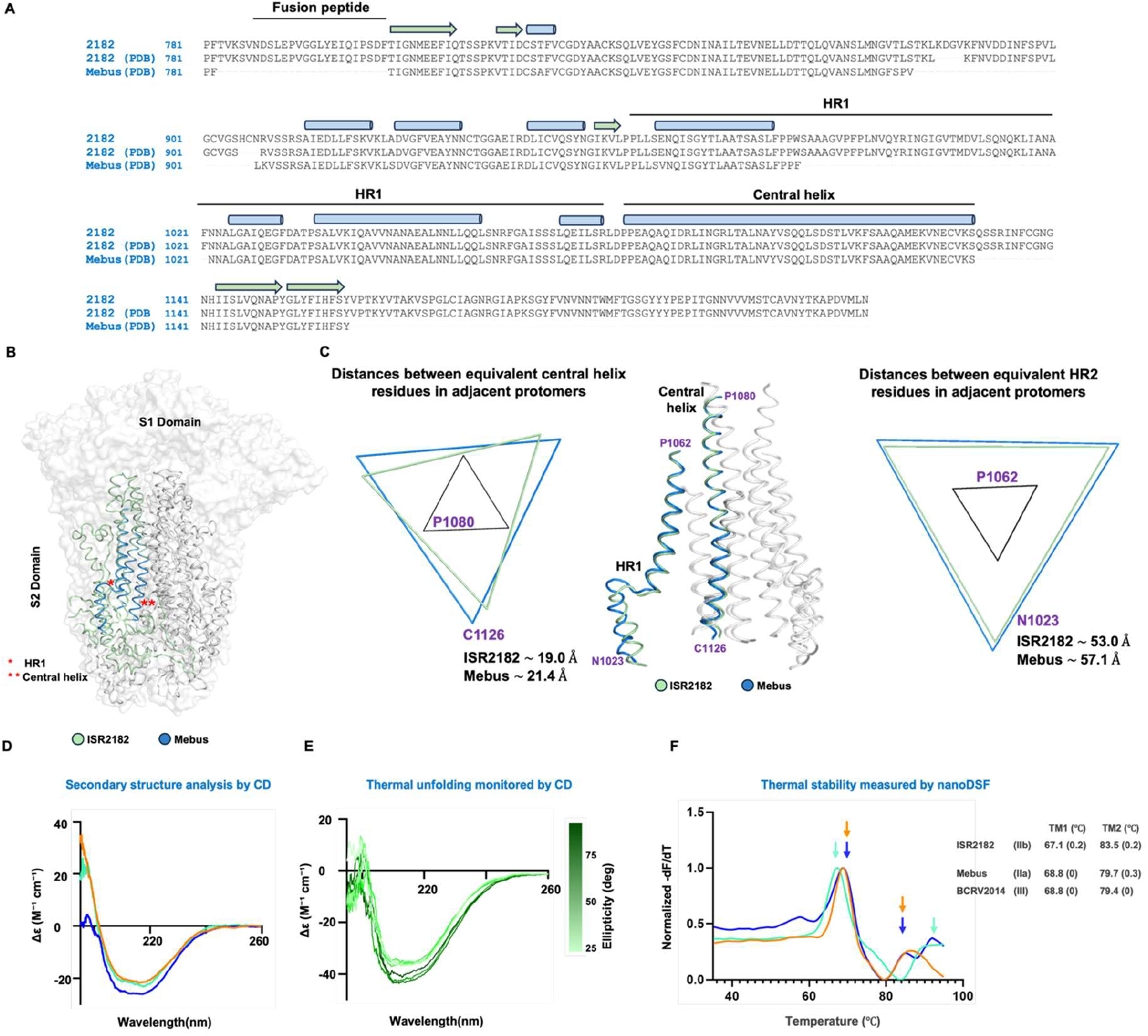
Structural basis and biophysical characterization of the Mebus spike stalk. **(A)** Sequence alignment of the S2 subunit from the BCoV Mebus and ISR2182 spike proteins highlighting the fusion peptide (FP), heptad repeat 1 (HR1), central helix (CH), and the membrane-proximal stalk region. Residues that are unresolved in the Mebus cryo-EM reconstruction originate from multiple discontinuous regions of the primary sequence that converge in the folded structure. Residues that are unresolved in the cryo-EM structures are indicated by dashes. **(B)** Superposition of the S2 subunits from ISR2182 and Mebus shown in cartoon presentation. One protomer is colored while the remaining two protomers are shown in gray. The HR1 and central helix are marked in red asterisks. The structures indicate that although the unresolved residues originate from discontinuous regions of the primary sequence, they cluster within the membrane-proximal stalk in the folded structure. **(C)** Superposition of the S2 helical bundles from ISR2182 (green) and Mebus (cyan). Distances between equivalent residues in adjacent protomers are shown for the central helix (P1080 and C1126) and HR1 (P1062 and N1023). Measurements of the distance between the residues at the head-proximal part (P1080; P1062) are similar in the two structures, whereas residues closer to the membrane-proximal stalk (C1126 and N1023) exhibit increased separation in the Mebus spike. **(D)** Far-UV CD spectra of purified ISR2182, Mebus, and BRCV2014 spike ectodomains showing similar overall secondary structure content. **(E)** Temperature-dependent CD measurements used to monitor thermal unfolding of the spike ectodomains. No well-defined cooperative melting transition was observed, preventing reliable determination of melting temperatures. **(F)** Thermal unfolding profiles of ISR2182, Mebus, and BRCV2014 spike ectodomains measured by nano differential scanning fluorimetry (nanoDSF). Arrows indicate the first (TM1) and second (TM2) melting transitions. Corresponding melting temperatures are summarized at right.

To investigate the structural basis underlying the increased conformational heterogeneity of the Mebus stalk, we compared the organization of the long α-helices that form the membrane-proximal S2 core, extending from the fusion peptide through heptad repeat 1 (HR1) and the central helix toward the membrane-proximal stalk (Fig. 3B). Superposition of the ISR2182 and Mebus structures showed that the helical bundle comprising HR1 and the central helix, adjacent to the S1 head, aligns closely between the two spikes. Consistent with this observation, the distances between equivalent residues in the adjacent protomers located within the upper portion of the bundle, including N1026 in HR1 and P1080 in the central helix, were identical in the two trimers (Fig. 3C). In contrast, residues located closer to the membrane-proximal region displayed substantially greater separation in the Mebus trimer. The distance between C1126 residues of the central helix in adjacent protomers increased from 19.0 Å in ISR2182 to 21.4 Å in Mebus, whereas the distance between N1023 residues in HR1 increased from 53.0 Å to 57.1 Å (Fig. 3C). Notably, no measurable differences were observed in the N-terminal helices adjacent to the fusion peptide, indicating that the structural divergence is restricted to the membrane-proximal portion of the helical bundle. These measurements indicate that the lower portion of the helical bundle adopts a more expanded configuration in the Mebus spike. Together, these observations indicate that the membrane-proximal portion of the S2 helical bundle adopts a more expanded configuration in Mebus, providing a structural explanation for the increased conformational heterogeneity of the downstream stalk.

### The increased conformational heterogeneity of the Mebus stalk is not associated with altered secondary structure

The absence of well-defined cryo-EM density could reflect either local unfolding or increased conformational heterogeneity. To distinguish between these possibilities, we compared the secondary structure content of the three spike proteins using circular dichroism (CD) spectroscopy (Fig. 3D). The CD spectra of ISR2182, Mebus, and BRCV2014 were nearly indistinguishable and exhibited the characteristic profile of CoV spike glycoproteins, consistent with a mixed α-helical and β-sheet secondary structure (Fig. 3D and Fig S3A) (Arruda *et al*, 2023). These findings indicate that the Mebus spike retains an overall secondary structure comparable to that of the contemporary BCoV lineages.

### Differential thermal stability among BCoV spike proteins

To determine whether the structural differences among the three BCoV spike proteins were associated with altered thermal stability, we first monitored thermal unfolding by CD spectroscopy. However, the resulting profiles lacked well-defined melting transitions, precluding reliable determination of melting temperatures (Fig. 3E). We therefore analyzed the spike ectodomains by nano differential scanning fluorimetry (nanoDSF). All three BCoV spike proteins exhibited biphasic thermal-unfolding profiles, consistent with previous analyses of coronavirus spike glycoproteins (Fig. 3F) (Lauster *et al*, 2025; Vollrath *et al*, 2014). The apparent transition temperatures nevertheless differed among the three strains.

ISR2182 displayed transitions at 67.3°C and 83.7°C, whereas Mebus and BRCV2014 exhibited nearly identical profiles, with transitions at 68.8°C and 79.4°C, and 68.8°C and 79.6°C, respectively.

Unexpectedly, the thermal unfolding profiles did not align with the structural differences observed by cryo-EM. Despite the pronounced conformational heterogeneity of its membrane-proximal stalk, the Mebus spike displayed thermal stability comparable to that of BRCV2014, whose stalk was well resolved. In contrast, ISR2182 exhibited a distinct unfolding profile, characterized primarily by a higher second transition temperature. These findings indicate that the increased conformational heterogeneity of the Mebus stalk does not arise from global destabilization of the spike trimer, but instead reflects a localized alternative conformational state.

### Antigenic conservation between human OC43 and bovine coronavirus spike proteins

The structural remodeling identified in the BCoV spike prompted us to ask whether these evolutionary changes have altered the antigenic landscape of the spike glycoprotein. This question is particularly relevant given the close evolutionary relationship between BCoV and the human coronavirus OC43, which shares approximately 91-92% amino acid identity with BCoV (Fig. S2A). Despite this high sequence conservation, sequence divergence is concentrated within the S1 subunit, particularly in the NTD and CTD, which contain the major neutralizing antibody epitopes. Consistent with these sequence differences, our structural analyses revealed localized remodeling of both domains (Figs. 2D-G and S2E-F). We therefore asked whether neutralizing antibodies raised against OC43 retain the ability to recognize the BCoV spike.

Previous structural studies defined the major neutralizing epitopes of the OC43 spike glycoprotein (Bangaru *et al*., 2022; Wang *et al*., 2022). These studies identified dominant antibody responses targeting both the NTD and CTD and determined cryo-EM structures of the OC43 spike bound to two representative neutralizing antibodies: 46c12, which recognizes the NTD, and 34E6, which binds the CTD (Fig. 4A). To predict whether these neutralizing antibodies could recognize BCoV, we superimposed the OC43 antibody-spike complexes onto the three BCoV spike structures. Structural analysis showed that 46c12 recognizes the receptor-binding L1 loop of the OC43 NTD, thereby sterically blocking receptor engagement (Fig. 4B). In contrast to the well-defined conformation observed in OC43, the corresponding L1 loop is unresolved in all three BCoV structures (Fig. S2E), indicating increased conformational heterogeneity. The absence of a defined receptor-binding loop in all three BCoV structures predicts that the 46c12 epitope cannot be formed in the same conformation as in OC43, suggesting that this antibody is unlikely to recognize the BCoV NTD (Fig. 4B). Similarly, structural analysis of the CTD-directed Ab 34E6 revealed that its epitope is centered on a surface-exposed loop spanning residues 460-480 (Fig. 4C). This loop exhibits relatively low sequence conservation between OC43 and BCoV and adopts a distinct conformation in the BCoV spike structures (Fig. 4D). Structural superposition predicted steric clashes between 34E6 and the remodeled BCoV loop, suggesting that these conformational differences would prevent Ab binding (Fig. 4C).

**Figure 4.**
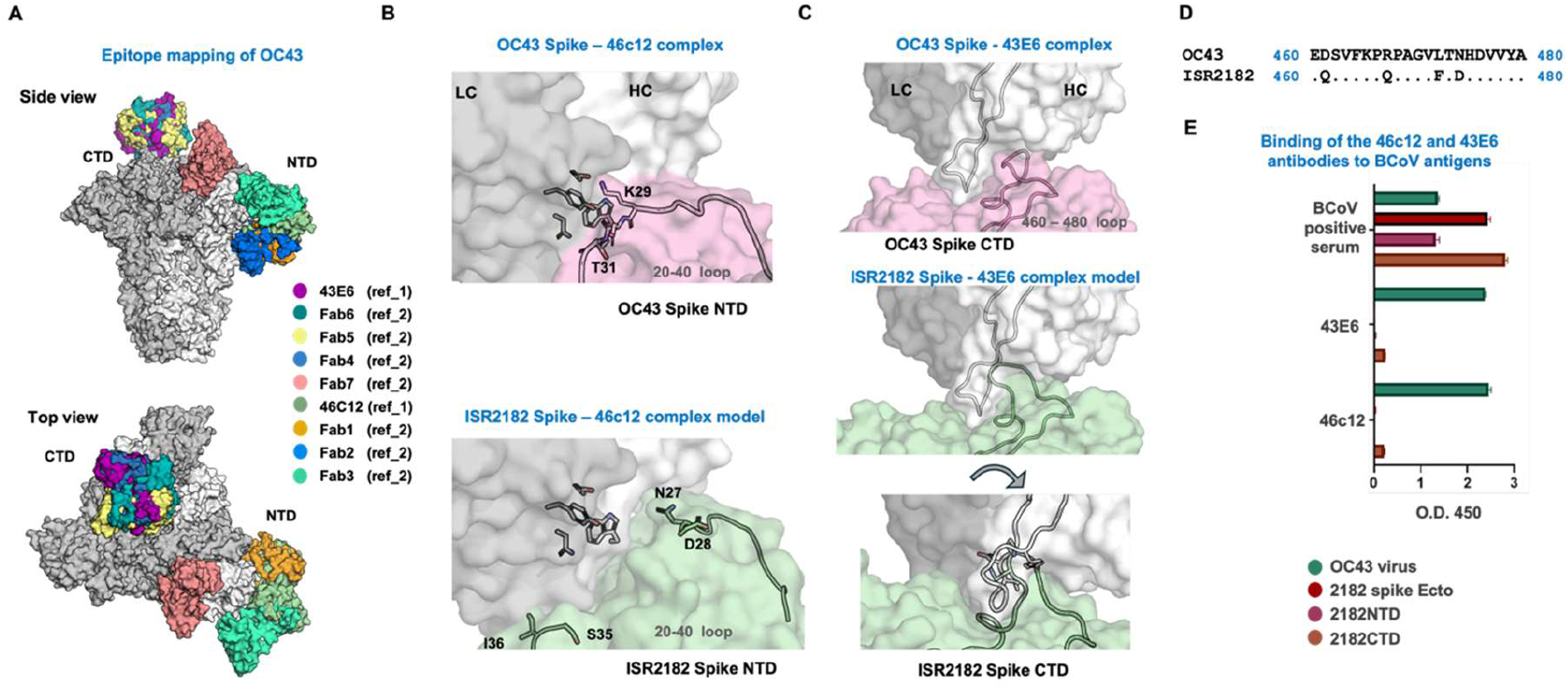
Structural basis for the antigenic divergence between OC43 and BCoV spike glycoproteins. **(A)** Antigenic landscape of the OC43 spike glycoprotein based on Cryo-EM structures of OC43-46C12 (PDB:7PNM) and OC43-43E6 (PDB:7PNQ) complexes and Cryo-EM polyclonal epitope mapping. For clarity, only the heavy- and light-chain variable regions are presented. **(B)** Structure of the OC43 spike in complex with the neutralizing antibody 46c12, which targets the NTD, showing recognition of the receptor-binding L1 loop (Top). Superposition of the 46c12-bound OC43 NTD onto the BCoV ISR2182 spike structure demonstrates that the unresolved L1 loop in BCoV is incompatible with the OC43 46C12 binding mode (Bottom). **(C)** Structure of the OC43 spike in complex with the neutralizing antibody 43E6, which targets the CTD, showing recognition of the surface-exposed 460-480 loop (Top). Superposition of the 43E6-bound OC43 CTD onto the corresponding region of the BCoV ISR2182 spike predicts steric clashes between the remodeled BCoV 460-480 loop and 43E6 (Bottom). **(D)** Sequence alignment of the 460-480 loop of OC43 and BCoV ISR2182. **(E)** Binding of OC43-specific neutralizing antibodies to 2182 spike. Recombinant ISR2182 NTD and CTD proteins were tested for binding to the OC43 neutralizing antibodies 46C12 and 43E6. Lysed OC43 virus served as a positive control. Both antibodies recognized the corresponding OC43 antigen but showed no detectable binding to the homologous BCoV domains, validating the structural predictions

### Experimental validation of the structural predictions

To experimentally validate the structural predictions, we expressed the NTD and CTD domains of ISR2182 and evaluated the binding of the neutralizing monoclonal Abs 46C12 and 34E6 by ELISA (Fig. 4D). As a control, we used lysed OC43 virus. Both 46c12 and 34E6 bound robustly to the corresponding OC43 antigens but exhibited no detectable binding to the homologous BCoV domains. These results validate the structural predictions and indicate that the antigenic differences between OC43 and BCoV are sufficient to prevent recognition by these representative OC43 antibodies. Together, these structural and biochemical analyses demonstrate that evolutionary remodeling of the BCoV spike has reshaped major neutralizing epitopes, resulting in substantial antigenic divergence from the closely related human coronavirus OC43 despite their high overall sequence conservation.

## Discussion

Our study demonstrates that BCoV evolution preserves the overall prefusion architecture of the spike glycoprotein while selectively remodeling key functional regions governing glycosylation, receptor recognition, conformational dynamics, and antigenicity. By integrating comparative genomics, glycoproteomics, cryo-electron microscopy, biophysical analyses, and antigenic characterization, we provide the first comparative structural analysis of representative BCoV lineages and establish a framework for understanding how evolution reshapes the structure and function of *Embecovirus* spike proteins.

One of the most striking examples of this remodeling is the emergence of a previously unrecognized N-linked glycosylation site at residue N1260, which is conserved among contemporary European BCoV isolates. Evolution of N-linked glycosylation has emerged as an important mechanism by which CoVs adapt to new hosts and evade immune recognition. In SARS-CoV-2, the acquisition or loss of individual glycans has been associated with altered antigenicity, immune evasion, and spike stability, whereas glycan remodeling has also been reported during the evolution of other *Betacoronaviruses* (Allen *et al*., 2023). Glycans within the S2 subunit are generally highly conserved across *Betacoronaviruses* because they contribute to protein folding, stabilization of the prefusion conformation, and membrane fusion (Allen *et al*., 2023; Watanabe *et al*., 2020). The identification of a lineage-specific glycan in BCoV therefore suggests that similar evolutionary pressures continue to shape *Embecovirus* spike proteins. Although the membrane-proximal location of N1260 prevented direct visualization by cryo-EM, glycoproteomic analysis confirmed that this site is occupied in the mature spike protein. Whether this lineage-specific glycan modulates spike stability, conformational dynamics, membrane fusion, or immune evasion remains an important question for future investigation.

Our analyses further demonstrate that BCoV evolution has remodeled the conformational landscape of the spike glycoprotein. Comparison with the closely related human coronavirus OC43 revealed pronounced flexibility of the BCoV receptor-binding L1 loop, whereas the equivalent region of OC43 adopts a well-defined preformed conformation (Tortorici *et al*., 2019). Interestingly, a recent structural study of contemporary OC43 isolates (Hassan *et al*, 2026) similarly demonstrated that ongoing evolution remodels the carbohydrate-binding site, resulting in altered receptor recognition and ligand-induced conformational changes. Together, these findings suggest that the receptor-binding surface of *Embecovirus* spike proteins remains evolutionarily plastic despite conservation of receptor usage. Such conformational plasticity may provide a mechanism to optimize receptor interactions while maintaining the overall spike architecture during viral evolution.

A second example of this structural remodeling was observed within the membrane-proximal stalk. Although the Mebus spike exhibited pronounced conformational heterogeneity in this region, secondary-structure analysis demonstrated that the stalk remained folded, indicating that the missing cryo-EM density reflects increased conformational sampling rather than structural unfolding.

Unexpectedly, these localized structural differences were not accompanied by reduced thermal stability. Instead, nanoDSF revealed lineage-specific thermal unfolding profiles that did not correlate with the degree of structural heterogeneity observed by cryo-EM. These findings suggest that local conformational dynamics and global thermodynamic stability evolve independently and emphasize that static structures alone may not fully capture the energetic landscape of coronavirus spike proteins.

The structural remodeling identified in this study also had important antigenic consequences. Despite sharing more than 90% amino acid sequence identity, BCoV and OC43 differed sufficiently in the NTD and CTD to abolish recognition by representative OC43-neutralizing antibodies (Bangaru *et al*., 2022; Wang *et al*., 2022). Structural modeling accurately predicted the loss of antibody binding, which was subsequently confirmed experimentally. These observations demonstrate that relatively localized structural remodeling can profoundly alter antigenic properties despite extensive overall sequence conservation. Given the proposed bovine origin of OC43, these findings further suggest that high sequence similarity alone should not be assumed to confer cross-protective immunity among closely related *Embecoviruses*.

Collectively, our findings support a model in which BCoV evolution proceeds through selective remodeling of key functional regions while preserving the overall architecture of the spike glycoprotein. Rather than accumulating neutral sequence variation alone, evolution has modified glycosylation, receptor-binding dynamics, conformational behavior, and antigenic surfaces in ways that are likely to influence viral fitness and host adaptation. Beyond providing the first structural comparison of representative BCoV lineages, this work establishes a framework for understanding *Embecovirus* spike evolution and highlights the importance of incorporating contemporary circulating strains into future structural studies, vaccine development, and surveillance efforts.

## Materials and Methods

### Samples collection and preparation

The clinical samples used in this study were selected from a pool of samples routinely submitted to the Kimron Veterinary Institute for BCoV diagnosis following suspicion of related disease. These samples were collected from sick and dead cattle (including calves, heifers, and dams) between 2017 and 2022. They included nasal and rectal swabs, as well as intestinal and lung tissues. These samples were processed as previously described (David *et al*., 2021).

### Molecular detection of BCoV from clinical samples

The presence of BCoV RNA in the clinical samples was confirmed by quantitative reverse transcriptase-polymerase chain reaction (RT-qPCR) using a fluorescent probe and a pair of primers targeting the BCoV nucleocapsid gene, as described previously (David *et al*., 2021; Kishimoto *et al*, 2017). Briefly, 50 µL of RNA was extracted from 100 µL of sample supernatant using a Ribospin^TM^ vRD II viral RNA purification kit (GeneAll Biotechnology, Co., ltd, Seoul, Korea) according to the manufacturer’s instructions. Five microliters of the purified RNA was immediately used to make an RT-qPCR reaction mix using Quanta qScript XLT 1-Step RT-qPCR ToughMix kit (Qunatabio, Beverly, MA, USA), according to the manufacturer’s instructions. The reaction tubes were then incubated and analyzed in the Bio-Rad CFX 96 Real-Time Detection System (Bio-Rad, Hercules, CA, USA).

### PCR and DNA preparation for sequencing

The complete spike and HE genes were amplified from the total RNA of BCoV-positive samples using Quanta qScript XLT 1-Step RT-PCR Kit (Qunatabio, Beverly, MA, USA). The PCR for the spike genes targeting the coding sequence (∼4092 bp) was performed using two independent primer pairs designed by us from the published nucleotide sequence of BCoV, the Mebus strain (GenBank accession No. BCU00735). Two primer pairs were used to amplify the complete spike coding sequence (pair 1: Forward_5’-ATGTTTTTGATACTTTTAATTTCC-3’ and Reverse_5’- TTAGTCGTCATGTGATGTTT-3’; pair 2: Forward_5’-GGTGGATAATGGTACTAGGCTGC-3’ and Reverse_5’-TCGTCAGGAGCCAATAAATCAA-3’). The PCR for the HE gene coding sequence was performed using the previously published primers (Park *et al*, 2006).

Following RT-PCR, the amplicons were verified by electrophoresis on a 1% agarose gel; the clear bands were excised and purified using a MEGAquick Spin Plus DNA fragment purification kit (iNTRON, Seoul, Korea). Purified PCR products were sent for sequencing.

### Sequencing of the complete spike and HE genes

For the spike genes, the PCR products were vacuum-dried, and concentrated DNA was sent for next-generation sequencing (NGS; Genotypic Technology, India). The DNA Libraries were prepared using the QIASeq FX DNA Library Preparation kit (Qiagen GmbH, Hilden, Germany), following the manufacturer’s instructions. Briefly, 10 to 50 ng of DNA was enzymatically fragmented, end-repaired, and A-tailed in a one-tube reaction using the FX Enzyme Mix. Next, the fragmented DNA was ligated to an Illumina adapter to generate a sequencing library, which was then amplified by Indexing-PCR to enrich adapter-tagged fragments. Finally, the amplified libraries were purified using a JetSeq clean-up kit (Bioline Reagents Ltd), quantified using a Qubit fluorometer (Thermo Fisher Scientific, MA, USA), and the fragment size distribution was analyzed on an Agilent 2200 TapeStation. The purified fragments were subjected to Illumina sequencing.

For the HE gene, the sequencing was performed using Sanger sequencing technology (Hylabs, Rehovot, Israel).

### Molecular analysis of the spike and HE protein sequences

For the spike gene, the Illumina short reads were *de novo* assembled in Geneious Prime 2022.2.2 (Biomatters, CA, USA). The consensus sequences of the contigs were assembled to reconstruct the full spike gene sequence. For the HE gene, the forward and reverse Sanger chromatograms were trimmed to remove low-quality ends, and the cleaned DNA sequences were *de novo* assembled in Geneious Prime 2022.2.2 (Biomatters, CA, USA).

### Phylogenetic analysis

For global molecular analysis, the spike and HE amino acid sequences of the reference BCoV strains were searched and downloaded from the NCBI database using BlastX. All the amino acid sequences were aligned using the Muscle sequence alignment tool in MEGA12.1 software (Kumar *et al*, 2024). The aligned sequences were analyzed for divergence in RStudio (RStudio, Posit PBC) to identify amino acid differences among BCoV strains and potential glycosylation sites on the spike and HE proteins. Phylogenetic analysis of the complete BCoV spike and HE amino acid sequences was performed using the neighbor-joining method in MEGA12.1 software (Kumar *et al*., 2024). The neighbor-joining trees were constructed using the Jones-Taylor-Thornton (JTT) amino acid substitution model with gamma-distributed rate variation among sites, and branch support was assessed using 1000 bootstrap replicates. The phylogenetic trees were visualized and annotated using Interactive Tree of Life (ITOL) (Letunic & Bork, 2024), and the final figure adjustments were performed in Inkscape 1.4.2 (Inkscape project, 2026, Inkscape).

### Plasmids Construction

Codon-optimized genes encoding prefusion-stabilized spike ectodomains (residues 15–1294) from Mebus, ISR2182, and BRCV2014 were synthesized (Twist Bioscience) and cloned into pDEST. Constructs contained a mutated furin cleavage site, five stabilizing proline substitutions, a C-terminal T4 fibritin foldon trimerization domain, a TEV cleavage site, an octa-histidine tag, and a tandem Strep-tag. The ISR2182 NTD and CTD plasmids were constructed from the ISR2182 Ecto plasmid by PCR followed by Gibson assembly.

### Expression and purification of stabilized BCoV spike ectodomains and ISR2182 NTD and CTD

Recombinant prefusion-stabilized spike ectodomains, ISR2182 NTD, and CTD were transiently expressed in Expi293F or Expi293S cells (Thermo Fisher Scientific) according to the manufacturer’s protocol. Culture supernatants were harvested five days post-transfection, clarified by centrifugation and filtration, and purified by Strep-Tactin affinity chromatography (Cytiva). Bound proteins were eluted with PBS supplemented with 2.5 mM desthiobiotin, concentrated, and further purified by size-exclusion chromatography on a Superdex 200 Increase 10/300 GL column (Cytiva) equilibrated in 20 mM Tris-HCl (pH 7.5) and 100 mM NaCl. Fractions corresponding to trimeric spike protein were concentrated, flash frozen, and used for cryo-EM, glycoproteomic, and biophysical analyses.

### Cloning and Purification of Antibodies

Variable regions of the heavy and light chains of the OC43 neutralizing antibodies 46C12 and 34E6 were synthesized (Twist Bioscience) and cloned into the mammalian expression vector pCMVtpa by Gibson Assembly. Recombinant antibodies were transiently expressed in Expi293 GnTI⁻ cells and purified from culture supernatants by Protein G affinity chromatography followed by size-exclusion chromatography on a Superdex 200 Increase 10/300 GL column (Cytiva). Antibody purity was confirmed by reducing and non-reducing SDS-PAGE.

### Sample preparation for glycoproteomic analysis

Purified ISR2182 spike protein expressed in Expi293S cells was treated with Endoglycosidase H (Endo H) to remove high-mannose and hybrid N-linked glycans while retaining a single N-acetylglucosamine residue on occupied glycosylation sites. Proteins were subsequently reduced with dithiothreitol, alkylated with iodoacetamide, and processed using S-Trap Micro Spin Columns (ProtiFi). On-column digestion was performed independently with trypsin or chymotrypsin to maximize peptide coverage. Peptides were desalted using homemade C18 StageTips and analyzed by LC-MS/MS. For more details, see Supplementary Methods.

### LC-MS/MS analysis

Peptide samples were analyzed by nanoLC-MS/MS using an Easy-nLC system coupled to a Q Exactive HF mass spectrometer (Thermo Fisher Scientific). Peptides were separated on a 25-cm EasySpray analytical column (45°C) using a 45-min linear gradient from 0% to 40% acetonitrile. Data were acquired in data-dependent acquisition (DDA) mode with full MS scans collected at 240,000 resolution (m/z 375–1,800), followed by HCD fragmentation of the 12 most abundant precursor ions (normalized collision energy, 26). MS/MS spectra were acquired at a resolution of 35,000 with dynamic exclusion set to 45 s. Additional acquisition parameters are provided in the Supplementary Methods.

### Glycoproteomic data analysis

Raw MS/MS data were analyzed using FragPipe (v23.1) with MSFragger against a custom protein database comprising the 100 most abundant proteins identified in the sample together with common contaminants and a reversed decoy database. Peptide and protein identifications were filtered to a 1% false discovery rate (FDR). To identify N-linked glycosylation sites following Endo H treatment, a variable modification corresponding to a residual N-acetylglucosamine (HexNAc; +203.07937 Da) on asparagine residues was included in the database search. Trypsin- and chymotrypsin-digested samples were searched using the appropriate enzyme specificity. Peptide quantification was performed using IonQuant, and Endo H-modified glycopeptides identified in both tryptic and chymotryptic digests were used to assign occupied N-linked glycosylation sites.

### Cryo-electron microscopy

Purified spike ectodomains from ISR2182, Mebus, and BRCV2014 were vitrified on glow-discharged Quantifoil R1.2/1.3 300-mesh copper grids and imaged using a Glacios transmission electron microscope (Thermo Fisher Scientific) operated at 200 kV and equipped with a Falcon 4i direct electron detector and a Selectris X energy filter. Movies were recorded in dose-fractionated counting mode using EPU (Thermo Fisher Scientific) at a calibrated pixel size of 0.89 Å, with a total electron dose of 30 e⁻/Å² and a defocus range of −0.7 to −2.0 μm.

Image processing was performed in CryoSPARC v4.3.0 (Punjani et al., 2017). Movies were corrected for beam-induced motion, and contrast transfer function (CTF) parameters were estimated using the patch-based algorithms implemented in CryoSPARC. Particles were template-picked, subjected to iterative two-dimensional (2D) classification, and used for ab initio reconstruction followed by three-dimensional (3D) refinement. Final maps were reconstructed to overall resolutions of 2.66 Å (ISR2182), 2.50 Å (Mebus), and 2.52 Å (BRCV2014), based on the gold-standard Fourier shell correlation criterion (FSC = 0.143). Detailed data collection and refinement statistics are provided in Supplementary Table 2. Additional information is provided in the Supplementary Methods.

### Nano differential scanning fluorimetry (nanoDSF)

The thermal stability of purified BCoV spike proteins was evaluated by nano differential scanning fluorimetry (nanoDSF) using a Prometheus instrument (NanoTemper Technologies). Purified proteins (0.6 mg/mL) were analyzed in buffer containing 20 mM Tris-HCl (pH 7.5) and 100 mM NaCl over a temperature range of 35–95°C. Melting temperatures (Tm) were determined from the first derivative of the ratio of intrinsic fluorescence measured at 350 and 330 nm (F350/F330) using the NanoTemper analysis software.

### Circular dichroism spectroscopy

Circular dichroism (CD) spectroscopy was performed to assess the secondary structure and thermal stability of purified BCoV spike ectodomains. Far-UV CD spectra were recorded between 190 and 260 nm, and thermal unfolding was monitored over a temperature range of 10–96°C. Spectra were normalized to molar ellipticity to enable comparison between samples analyzed at different protein concentrations. Detailed experimental procedures and data analysis are provided in the Supplementary Methods.

### ELISA to test binding of OC43 antibodies to BCoV spike

Half-area, clear, high-binding microlon microplates (Greiner Bio-One, Germany) were coated with 0.5 µg/mL of purified recombinant ISR2182 spike, NTD, or CTD protein and incubated overnight at 4 °C. Inactivated OC43 virus was used as a control. The plates were emptied and washed 3 times with PBS, then blocked with 10% seronegative fetal bovine serum (FBS) for 1 hour. The plates were washed 5 times with phosphate-buffered saline containing 0.05% Tween-20 (PBST 0.05%) and incubated with OC43 Abs at 10 µg/mL for 1 hour at room temperature. BCoV positive serum (at 1:100) was used as a control. Goat anti-human IgG FC HRP (1:2000) was added after 6 washes and incubated for another hour at room temperature. For the positive serum, goat anti-bovine HRP (1:50,000) was used. The plates were washed 5 times, incubated for 20 min in the dark with TMB substrate (Biolegend, San Diego, USA), and the reaction was stopped with 2 M sulphuric acid (Merck KGaA, Darmstadt, Germany). The absorbance of the reaction colors was measured at 450 nm in a Synergy HI microplate reader and analyzed using Gen5 version 3.09 software (BioTek Instruments, Inc., Vermont, USA).

## Acknowledgements

This work is funded in part by the Israel Dairy Board Foundation, grants # 828-0001 (to N.T.) and #848- 0349 (to A.S.).

## Author contributions

A.S and N.T. conceived and designed the study; J.A. and A.S. isolated the Israeli BCoV strain and sequenced the Spike and HE genes; J.A., A.S., and N.T. performed the phylogenetic analysis; H.T. and J.A. expressed and purified the recombinant proteins; M.S., N.K., H.T., and N.T. performed the glycoproteomic studies; Y.L.K., A.S. and R.Z. performed the cryo-EM data collection and processing; H.T. and N.T. solved and analyzed the structures; H.T. performed CD and nanoDSF experiments; J.A., H.T., A.S, and N.T. analyzed the data and wrote the manuscript with input from all authors. All authors reviewed and approved the final manuscript.

## Data availability

All data supporting the findings of this study are available within the paper and its Supplementary Information. Additional data are available from the corresponding authors upon reasonable request. The mass spectrometry proteomics data have been deposited to the ProteomeXchange Consortium via the PRIDE (Perez-Riverol *et al*, 2025) partner repository with the dataset identifier PXD081716.

Reviewers can access the PRIDE website using the following details: Project accession: PXD081716 Token: 5XfsyZSIj3n2

